# Expression of Spred2 in the urothelial tumorigenesis of the urinary bladder

**DOI:** 10.1101/2021.07.23.453537

**Authors:** Shinsuke Oda, Masayoshi Fujisawa, Li Chunning, Toshihiro Ito, Takahiro Yamaguchi, Teizo Yoshimura, Akihiro Matsukawa

## Abstract

Aberrant activation of the Ras/Raf/ERK (extracellular-signal-regulated kinase)-MAPK (mitogen-activated protein kinase) pathway is involved in the progression of cancer, including urothelial carcinoma; but the negative regulation remains unclear. In the present study, we investigated pathological expression of Spred2 (Sprouty-related EVH1 domain-containing protein 2), a negative regulator of the Ras/Raf/ERK-MAPK pathway, and the relation to ERK activation and Ki67 index in various categories of 275 urothelial tumors obtained from clinical patients. In situ hybridization demonstrated that Spred2 mRNA was highly expressed in high-grade non-invasive papillary urothelial carcinoma (HGPUC), and the expression was decreased in carcinoma in situ (CIS) and infiltrating urothelial carcinoma (IUC). Immunohistochemically, membranous Spred2 expression, important to interact with Ras/Raf, was preferentially found in HGPUC. Interestingly, membranous Spred2 expression was decreased in CIS and IUC relative to HGPUC, while ERK activation and the expression of the cell proliferation marker Ki67 index were increased. HGPUC with membranous Spred2 expression correlated significantly with lower levels of ERK activation and Ki67 index as compared to those with negative Spred2 expression. Thus, our pathological findings suggest that Spred2 negatively regulates cancer progression in non-invasive papillary carcinoma possibly through inhibiting the Ras/Raf/ERK-MAPK pathway, but this regulatory mechanism is lost in cancers with high malignancy. Spred2 appears to be a key regulator in the progression of non-invasive bladder carcinoma.

## Introduction

Bladder cancer is a highly prevalent disease and its incidence is steadily rising worldwide [1]. In the United States, bladder cancer is the 4th most incident and 8th most deadly tumor among men [2]. Most of the bladder cancer is urothelial carcinoma arising from urothelial epithelium. Evidence indicates that urothelial carcinoma has two distinct clinical subtypes with distinct molecular features at bladder tumor initiation; low-grade tumors (superficial papillary) and high-grade tumors (flat, represented by carcinoma in situ) [3, 4]. Low-grade tumors, i.e., papillary urothelial neoplasm of low malignant potential or low-grade papillary urothelial carcinoma, do not easily progress to high-grade papillary urothelial carcinoma or invasive carcinoma [5, 6]. Recently, a comprehensive landscape of molecular alterations in urothelial carcinomas was shown [7]. More than 70% of low-grade papillary carcinomas harbor FGFR3 gene mutation [8]. On the other hand, flat carcinoma in situ (CIS) often develops to invasive urothelial carcinoma [9, 10], in which allelic deletion of the *p53* and *PTEN* (tumor-suppressor) [11] and *retinoblastoma* gene (RB, negative cell cycle regulator) [12] is common.

In addition to the gain of function gene mutations, extracellular-regulated kinase (ERK) plays a crucial role in cancer development and progression [13, 14]. The Ras/Raf/ERK-MAPK (mitogen-activated protein kinase) pathway, one of the serine/threonine kinases of MAPKs pathway, is a major determinant to promote cell proliferation, differentiation, and survival, and plays an important role in bladder cancer prognosis [15]. ERK activation was observed in high-grade non-invasive and invasive urothelial carcinoma [16], suggesting that robust ERK activation contributes to urothelial tumorigenesis with a high malignant potential.

Signaling pathways are counterbalanced by endogenous inhibitory mechanism(s). Spred2 (Sprouty-related, EVH1 domain-containing protein 2) inhibits Ras-dependent ERK signaling by suppressing the phosphorylation and activation of Raf [17]. Ras activation is aberrant in many tumors due to oncogenic mutation of the *Ras* genes or alterations in upstream signaling components [18]. Rational therapies that target the Ras/Raf/ERK-MAPK pathway continues to attract much attention for cancer therapy [19].

We have hitherto investigated in different types of murine models and found that Spred2 controls inflammation by down-regulating the Ras/Raf/ERK-MAPK pathway [20–29]. Interestingly, Spred2 expression is down-regulated in invasive carcinomas such as hepatocellular carcinoma [30, 31] and prostatic adenocarcinoma [32]. Thus, altered Spred2 expression could affect urothelial tumorigenesis by regulating the Ras/Raf/ERK-MAPK pathway in bladder cancer. However; the pathophysiological roles of Spred2 in bladder cancer tumorigenesis remain largely unknown. In the present study, we examined the mRNA and protein expression of Spred2 in a range of human urothelial tumors. Our present findings suggest that endogenous Spred2 affects urothelial cancer progression, especially in non-invasive papillary urothelial carcinoma.

## Materials and methods

### Clinical samples

A total of 275 bladder biopsy or resection specimens (transurethral resection and cystectomy) during the year 2001-2016 were retrieved from pathology record at Department of Pathology, Okayama University Hospital. The patients who underwent chemotherapy or radiotherapy before the resection were not included in this study. All the hematoxylin and eosin (HE)-stained sections were reviewed and categorized by two blinded pathologists according to the 2016 WHO classification: non-tumor urothelium (non-tumor), papillary urothelial neoplasm of low malignant potential (PUNLMP), low-grade papillary urothelial carcinoma (LGPUC), high-grade papillary urothelial carcinoma (HGPUC), carcinoma in situ (CIS), and infiltrating urothelial carcinoma (IUC). All sections were used for immunohistochemistry. For in situ hybridization, sections were randomly chosen from each category. Cases for the enrolled 275 patients were shown in Table 1, in which clinicopathological features of each category were noted.

**Table 1.** Cases for the enrolled 275 patients.

|  | cases (%) | number of examined<br>by IHC/ISH | features |  |  |
| --- | --- | --- | --- | --- | --- |
|  |  |  | progression | nuclear grade | invasiveness |
| non-tumor | 101 (36.7) | 101/10 | - | - | - |
| PUNLMP | 19 (6.9) | 19/10 | slow | very low | non-invasive |
| LGPUC | 41 (14.9) | 41/15 | slow | low | non-invasive |
| HGPUC | 43 (15.6) | 43/18 | quick | high | non-invasive |
| CIS | 39 (14.2) | 39/18 | quick | high | non-invasive |
| IUC | 32 (11.6) | 32/14 | quick | high>>low | invasive |

non-tumor; non-tumor urothelium, PUNLMP; papillary urothelial neoplasm of low malignant potential, LGPUC; low-grade papillary urothelial carcinoma, HGPUC; high-grade urothelial carcinoma, CIS; carcinoma in situ, IUC; infiltrating urothelial carcinoma. IHC; immunohistochemistry, ISH; in situ hybridization.

The protocol in this study was reviewed and approved by the *Ethics Committee, Okayama University Graduate School of Medicine, Dentistry and Pharmaceutical Sciences and Okayama University Hospital (1608-009)*. Informed consent was obtained in the form of opt-out on our website. Those who rejected were excluded. This consent procedure conformed to amended Ethical Guidelines for Clinical Studies provided by Ministry of Health, Labor and Welfare of Japan (May 31, 2015) was approved by the *Ethics Committee, Okayama University Graduate School of Medicine, Dentistry and Pharmaceutical Sciences and Okayama University Hospital*.

### In situ hybridization

A total of 85 samples were randomly selected from 275 samples (Table 1). Paraffin-embedded tissue samples were sectioned at 5-μm-thick, kept on glass slides overnight at 45°C and then in situ hybridization was performed using the Affymetrix ViewRNA ISH Tissue Assay kit (QVT0050) and ViewRNA Chromogenic Signal Amplification kit (QVT0200) (Thermo Fisher Scientific, MA, USA), according to the manufacturer’s instructions. Human Spred2 probe set was purchased from Thermo Fisher (Affymetrix, Catalog No. VA1-17417-01). Spred2 mRNA expression was stained in red-dot. The total number of red-dot in 100 cells was counted in each sample under microscope by two blinded pathologists, and the number of red-dot per cell was calculated.

### Immunohistochemistry

For immunohistochemistry, all 275 specimens were employed (Table 1). Immunostaining for Spred2 was carried out using the Polink-2 plus HRP rabbit with DAB kit (GBI, Bothell, WA, USA), according to the manufacturer’s instructions. In brief, sections (4-µ-thick) were treated by microwave oven in 0.1 M citric acid buffer, treated with 3%H_2_O_2_ in methanol, blocked with DAKO Protein Block Serum-Free (Dako, Carpinteria, CA, USA), and incubated with anti-human Spred2 polyclonal antibody (Proteintech, Rosemont, IL, USA). Sections were then incubated with rabbit antibody specific enhancer, followed by the addition of polymer-HRP for rabbit IgG, and visualized using DAB complex. Nuclear counterstaining was performed using hematoxylin. Immunostaining for pERK1/2 (Clone D13.14.4E, Cell Signaling Technology, Danvers, MA, USA) and Ki67 (Clone MIB-1, Dako) was performed on a Ventana Discovery XT automated stainer (Ventana, Tucson, AZ, USA) with using iVIEW DAB Detection Kit (Ventana).

### Evaluation of immunohistochemistry

Spred2 was stained in the cytoplasm (C) or/and membrane (M). Immunoreactivity for Spred2 was classified into 4 groups, according to subcellular localization and staining intensity; C-M-, absent or weak staining intensity in cytoplasm and membrane; C-M+, moderate to strong membranous staining without staining in cytoplasm; C+M-, moderate to strong cytoplasmic staining without membranous staining; C+M+, moderate to strong cytoplasmic and membranous staining. pERK immunostaining was scored on the following semiquantitative scale as previously reported with modifications [33]: no staining (0); focal to <10% of cells (1); 10-50% of cells (2); 50% or more cells stained weak (3); 10-50% stained strong (4); 50% or more stained strong (5). Ki67 index, a marker for cell proliferation, was determined by counting 500 tumor cells, and the percentages of positively stained cells were determined. The stained sections were assessed by two blinded pathologists.

### Database analysis

Datasets with more than 25 samples in each category from Sanchez-Carbayo bladder 2 [34], Blaveri bladder 2 [35], and Stransky bladder [36] were used to analyze Spred2 expression in bladder cancer. An unpaired two-tailed t test was used for the statistical analysis. Kaplan-Meier Plotter (http://www.kmplot.com) was used to analyze the prognostic values of *Spred2* mRNA expression levels in bladder carcinoma. Kaplan-Meier survival plots were drawn using data from the Kaplan-Meier database. A log-rank *p*-value <0.05 was considered to indicate a statistically significant difference.

### Statistical analysis

Statistical analysis was performed using GraphPad Prism7 (GraphPad Software, San Diego, CA, USA) and js-STAR (free software). Dunn’s multiple comparison test was performed after Kruskal-Wallis test for the comparison of mean values among multi-groups. Multiple Fisher’s exact test was performed using the Bonferroni correction for the comparison of proportions among multi-groups. Mann-Whitney test was used for the comparison of mean values between the two groups. A value of *p*<0.05 was considered statistically significant.

## Results

### Spred2 mRNA expression in bladder urothelial tumors

We first examined Spred2 mRNA expression in various categories of 85 urothelial lesions including non-tumor, PUNLMP, LGPUC, HGPUC, CIS, and IUC. Figure 1A shows the representative HE and in situ hybridization photographs from each category, in which Spred2 mRNA expression was presented by red-dot (Fig 1A). The number of red-dots per cell was regarded as Spred2 mRNA expression level (Fig 1B). Levels of Spred2 mRNA expression were increased as the malignancy of the cancer increased in papillary tumors. Of note, the level reached the peak in HGPUC and then decreased in CIS and IUC. Spred2 mRNA expression in IUC was significantly lower than that in HGPUC (Fig 1B). These results indicate that Spred2 mRNA expression was up-regulated in non-invasive papillary bladder cancer as compared to cancers with high malignancy including invasive carcinoma.

**Fig 1.**
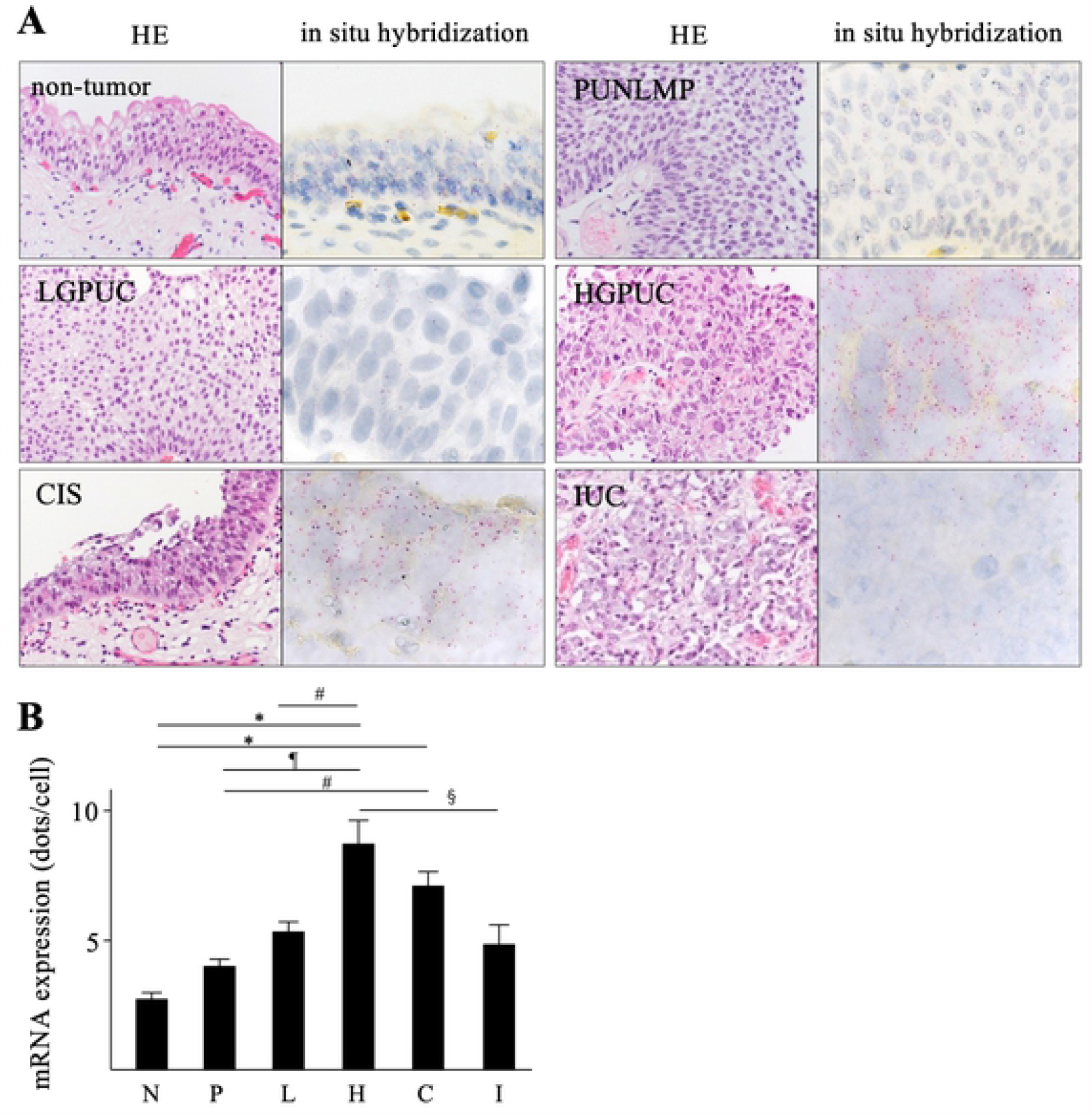
Spred2 mRNA expression in urothelial tumors. (A) Representative photographs of HE-(original magnification 400×) and in situ hybridization-sections from each category are shown. Spred2 mRNA expression was presented by red dots. (B) The number of the red-dots per cell was counted under microscope and Spred2 mRNA expression level was shown per one cell from each category (N: non-tumor; n=10, P: PUNLMP; n=10, L: LGPUC; n=15, H: HGPUC; n=18, C: CIS; n=18, and I: IUC; n=14). Data were mean ± SEM. ^#^*p*<0.05, ^§^*p*<0.01, ^¶^*p*<0.001, \**p*<0.0001 (Dunn’s multiple comparison test).

### Spred2 protein expression in bladder urothelial tumors

We next examined Spred2 protein expression by immunohistochemistry in 275 bladder urothelial tumors. To confirm immunoreactivity of Spred2 antibody, H1993 cells were stained with the antibody under overexpressing Spred2 (Supplementary Fig 1). Spred2 protein expression (Fig 2A) was immunophenotypically classified into 4 groups, according to the subcellular localization and staining intensity. The staining pattern in each tumor category was shown in Table 2. In all non-tumor cases, Spred2 was positive in cytoplasm of basal and lower intermediate cells (pattern C+M-, 101/101 cases). More than half of the cases of PUNLMP, CIS, and IUC showed absent or weak staining (C-M-; 74% (14/19 cases), 74% (29/39 cases), and 69% (22/32 cases), respectively). LGPUC and HGPUC showed membranous staining (C-M+ and C+M+) more frequently (49% (20/41 cases), and 51% (28/43 cases), respectively) than other categories (Table 2). We then compared mRNA and protein expression of Spred2. Cases with membranous staining, regardless of cytoplasmic staining pattern (C-M+ and C+M+), showed significantly higher levels of Spred2 mRNA expression than those without membranous staining (C-M- and C+M-) (Fig 2B). Spred2 is a membrane-associated substrate of receptor tyrosine kinase [17, 37] and react with Raf localized in the raft domain of the plasma membrane [38], suggesting that membranous Spred2 is more meaningful when considering the functional regulation. The positive rate of membranous Spred2 expression (C-M+ and C+M+) in each category was shown in Figure 2C, which showed that the expression was increased in LGPUC, peaked in HGPUC and then decreased in CIS and IUC. Together with the mRNA expression data, these results suggest that functional Spred2 was most expressed in HGPUC, and the expression was lower in CIS and IUC as compared to HGPUC.

**Table 2.** Subcellular immunolocalization of Spred2 in each tumor category

|  | C-M- | C+M- | C-M+ | C+M+ | total cases |
| --- | --- | --- | --- | --- | --- |
| non-tumor | 0 | 101 | 0 | 0 | 101 |
| PUNLMP | 14 | 4 | 1 | 0 | 19 |
| LGPUC | 15 | 6 | 20 | 0 | 41 |
| HGPUC | 8 | 7 | 22 | 6 | 43 |
| CIS | 29 | 4 | 4 | 2 | 39 |
| IUC | 22 | 0 | 7 | 3 | 32 |
| total cases | 88 | 122 | 54 | 11 | 275 |

non-tumor; non-tumor urothelium, PUNLMP; papillary urothelial neoplasm of low malignant potential, LGPUC; low-grade papillary urothelial carcinoma, HGPUC; high-grade urothelial carcinoma, CIS; carcinoma in situ, IUC; infiltrating urothelial carcinoma. IHC; immunohistochemistry, ISH; in situ hybridization. C; cytoplasm, M; membrane.

**Fig 2.**
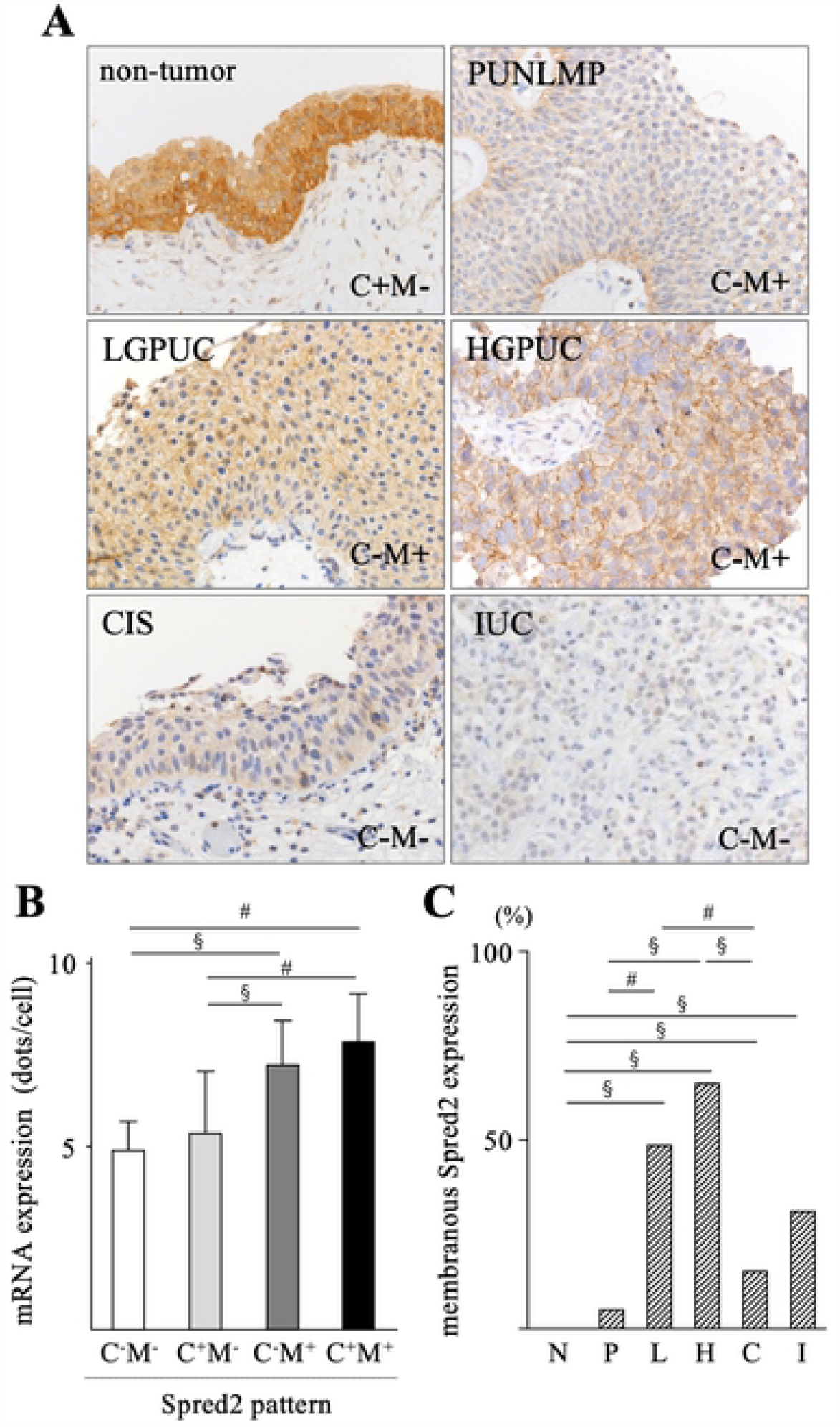
Immunohistochemical analyses of Spred2 protein expression in urothelial tumors. (A) Representative photographs of Spred2 immunohistochemistry (original magnification 400×) from each category are shown. (B) Expression levels of Spred2 mRNA in each Spred2 staining pattern were shown. C; cytoplasm, M; membrane. (C-M-; n=81, C+M-; n=122, C-M+; n=54, C+M+; n=11). Data were mean ± SEM. ^#^*p*<0.05, ^§^*p*<0.01 (Dunn’s multiple comparison test). (C) The positive rate of membranous Spred2 expression in each category was shown (N: non-tumor; n=101, P: PUNLMP; n=19, L: LGPUC; n=41, H: HGPUC; n=43, C: CIS; n=39, and I: IUC; n=32). Data were mean ± SEM. ^#^*p*<0.05, ^§^*p*<0.01 (Multiple Fisher’s exact test).

### Expression of pERK and Ki67 in bladder urothelial tumors

Increased Spred2 expression may affect the activation of the Ras/Raf/ERK-MAPK pathway and subsequent cancer growth. To address this possibility, we investigated the protein expression of phosphorylated ERK (pERK), an indicator of ERK-MAPK activation status, by immunohistochemically in each category. pERK was detected in the nucleus and cytoplasm of urothelial epithelial lesions in all specimens from each category with different intensity in strength (Fig 3A). The intensity of nuclear and cytoplasmic staining was then evaluated. Weak pERK staining (score, 1 and 2) was detected in 87% (score 1; 67/101, score 2; 21/101 cases) and 100% (score 1; 13/19, score 2; 6/19 cases) of non-tumor and PUNLMP, respectively (Table 3). In cancer categories (LGPUC, HGPUC, CIS and IUC), cancer cells with moderate (score 3) and strong (score 4 and 5) staining were increased. Strong pERK staining was detected in 10% (score 4; 4/41, score 5; 0/41), 44% (score 4; 14/43, score 5; 5/43 cases), 56% (score 4; 14/39, score 5; 8/39 cases), and 78% (score 4; 16/32, score 5; 9/32 cases) in LGPUC, HGPUC, CIS and IUC, respectively (Table 3). pERK score in each category was increased according to the malignant potential (Fig 3B). We next performed Ki67 immunohistochemistry (Fig 4A) and calculated Ki67 index (Fig 4B), an indicator of cell proliferation marker, in all categories. Ki67 index was significantly increased in all categories of bladder urothelial tumors as compared to non-tumor. Ki67 index of HGPUC, CIS and IUC was significantly higher than that of LGPUC (Fig 4B). Thus, pERK score and Ki67 index increase with high malignancy.

**Table 3.** pERK score in Spred2 immunostaining pattern

| Spred2 pattern \ pERK score |  | 1 | 2 | 3 | 4 | 5 | total cases |
| --- | --- | --- | --- | --- | --- | --- | --- |
| non-tumor | C-M- | 0 | 0 | 0 | 0 | 0 | 0 |
|  | C+M- | 67 | 21 | 13 | 0 | 0 | 101 |
|  | C-M+ | 0 | 0 | 0 | 0 | 0 | 0 |
|  | C+M+ | 0 | 0 | 0 | 0 | 0 | 0 |
| PUNLMP | C-M- | 11 | 3 | 0 | 0 | 0 | 14 |
|  | C+M- | 2 | 2 | 0 | 0 | 0 | 4 |
|  | C-M+ | 0 | 1 | 0 | 0 | 0 | 1 |
|  | C+M+ | 0 | 0 | 0 | 0 | 0 | 0 |
| LGPUC | C-M- | 3 | 4 | 8 | 0 | 0 | 15 |
|  | C+M- | 1 | 0 | 3 | 2 | 0 | 6 |
|  | C-M+ | 0 | 5 | 13 | 2 | 0 | 20 |
|  | C+M+ | 0 | 0 | 0 | 0 | 0 | 0 |
| HGPUC | C-M- | 0 | 1 | 2 | 4 | 1 | 8 |
|  | C+M- | 0 | 0 | 2 | 3 | 2 | 7 |
|  | C-M+ | 0 | 4 | 14 | 4 | 0 | 22 |
|  | C+M+ | 0 | 0 | 1 | 3 | 2 | 6 |
| CIS | C-M- | 0 | 1 | 11 | 11 | 6 | 29 |
|  | C+M- | 0 | 0 | 2 | 1 | 1 | 4 |
|  | C-M+ | 0 | 0 | 2 | 1 | 1 | 4 |
|  | C+M+ | 0 | 0 | 1 | 1 | 0 | 2 |
| IUC | C-M- | 0 | 0 | 4 | 11 | 7 | 22 |
|  | C+M- | 0 | 0 | 0 | 0 | 0 | 0 |
|  | C-M+ | 0 | 1 | 1 | 3 | 2 | 7 |
|  | C+M+ | 0 | 0 | 1 | 2 | 0 | 3 |

non-tumor; non-tumor urothelium, PUNLMP; papillary urothelial neoplasm of low malignant potential, LGPUC; low-grade papillary urothelial carcinoma, HGPUC; high-grade urothelial carcinoma, CIS; carcinoma in situ, IUC; infiltrating urothelial carcinoma. IHC; immunohistochemistry, ISH; in situ hybridization.

**Fig 3.**
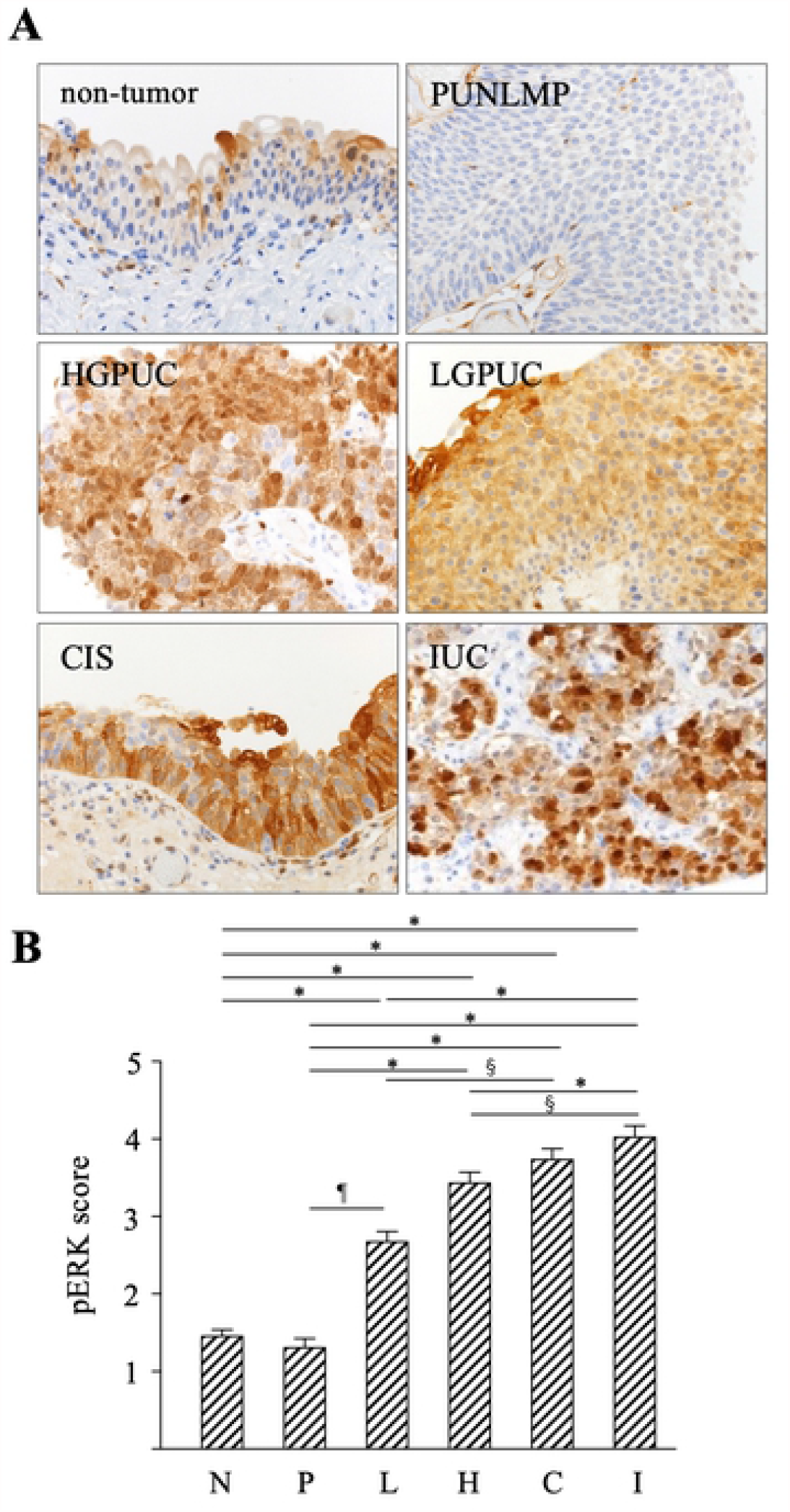
pERK score in urothelial tumors. (A) Representative photographs of pERK immunohistochemistry (original magnification 400×) from each category are shown. (B) pERK staining intensity was evaluated and scored (0-5), and pERK score in each category was shown (N: non-tumor; n=101, P: PUNLMP; n=19, L: LGPUC; n=41, H: HGPUC; n=43, C: CIS; n=39, and I: IUC; n=32). Data were mean ± SEM. ^§^*p*<0.01, ^¶^*p*<0.001, ^*^*p*<0.0001 (Dunn’s multiple comparison test).

**Fig 4.**
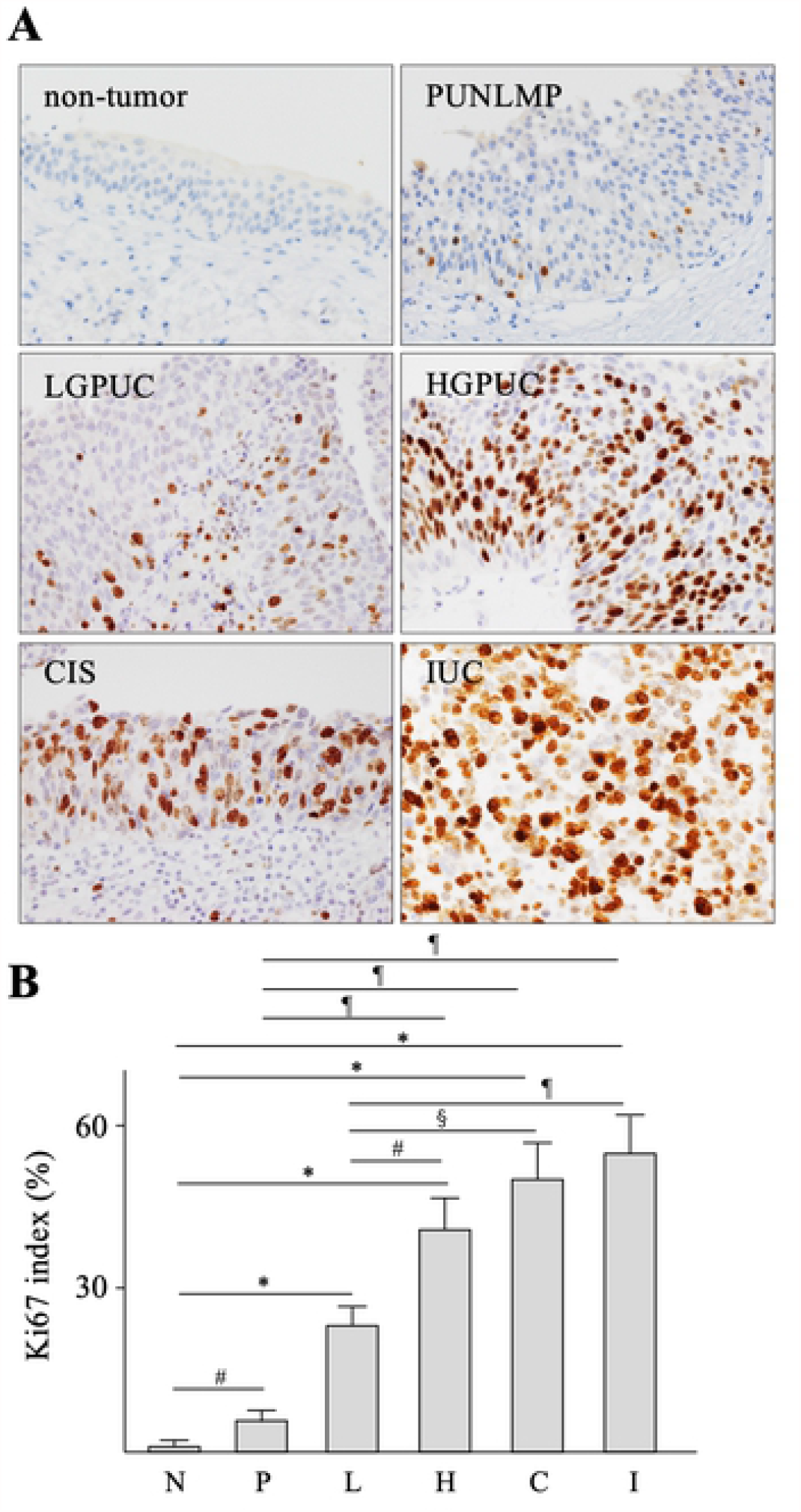
Ki67 index in urothelial tumors. (A) Representative photographs of Ki67 immunohistochemistry (original magnification 400×) from each category are shown. (B) Ki67 index in each category was shown (N: non-tumor; n=101, P: PUNLMP; n=19, L: LGPUC; n=41, H: HGPUC; n=43, C: CIS; n=39, and I: IUC; n=32). Data were mean ± SEM. ^#^*p*<0.05, ^§^*p*<0.01, ^¶^*p*<0.001, \**p*<0.0001 (Dunn’s multiple comparison test).

### Comparison between pERK score/Ki67 index and membranous Spred2 expression

We next compared the relation between pERK score/Ki67 index and membranous Spred2 expression (negative: M-, positive: M+) in cancer categories. In HGPUC, pERK score with Spred2 M+ were lower than those with Spred2 M-. No differences were found in LGPUC, CIS and IUC (Fig 5A). Since an increase in pERK is generally associated with an increased Ki67 index [39], ERK activation may result in increased tumor cell proliferation. As shown in Figure 5B, Ki67 index in HGPUC with Spred2 M+ was lower than those with Spred2 M-. These results suggest that membranous Spred2 plays a role in down-regulated ERK activation and subsequent cancer cell proliferation in HGPUC, but this negative regulatory mechanism is not functioning in CIS. Although pERK score was not different between Spred2 M- and Spred2 M+ in IUC, Ki67 index was decreased in Spred2 M+ as compared to Spred2 M-, indicating that Spred2 may downregulate cancer cell proliferation through ERK-MAPK independent pathway in IUC.

**Figure 5.**
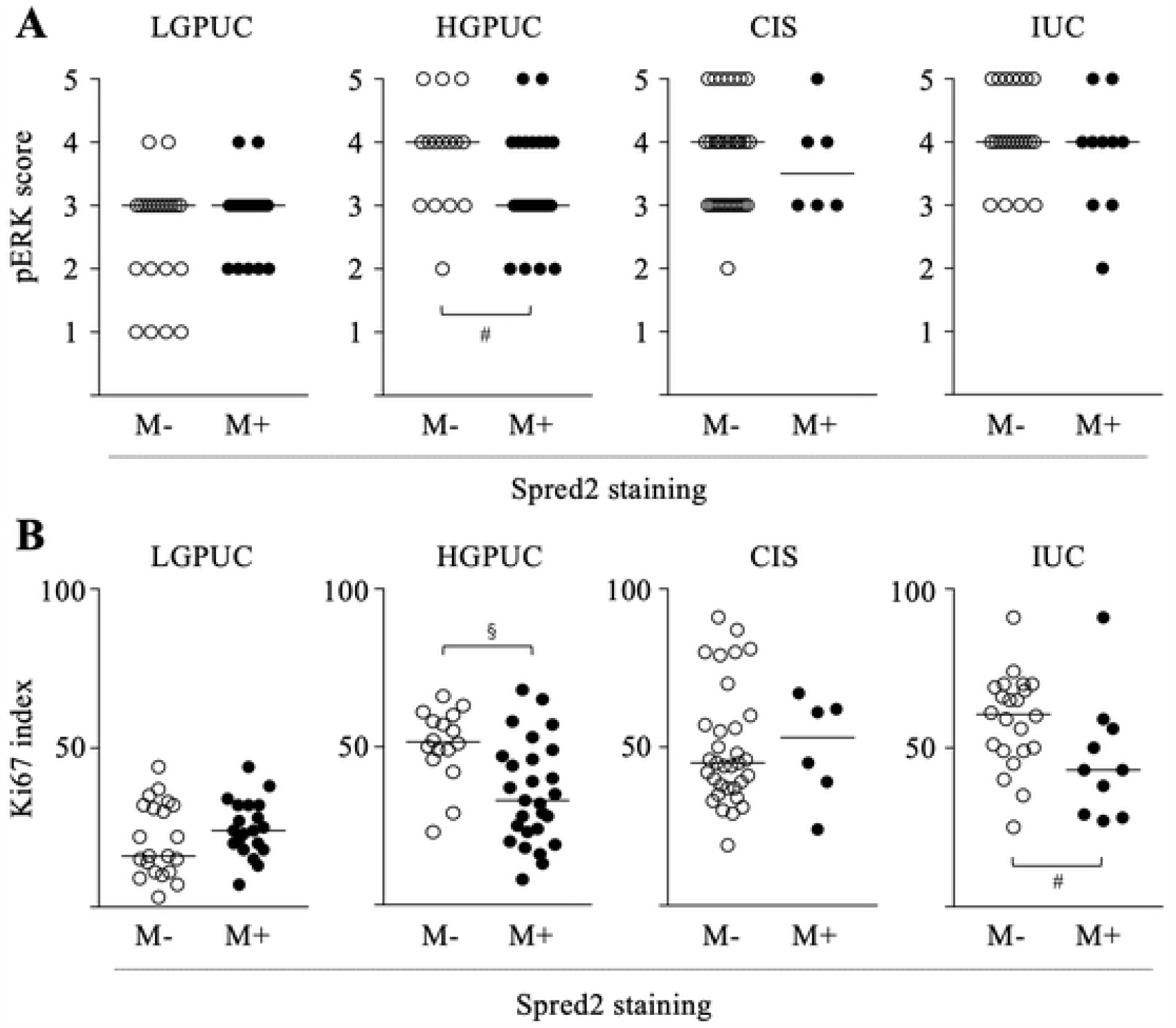
Comparison between pERK score/Ki67 index and membranous Spred2 expression. pERK score (A) and Ki67 index (B) were compared between membranous Spred2 negative (M-) and positive (M+) in cancer categories (LGPUC; n=41, HGPUC; n=43, CIS; n=39, and IUC; n=32). Bar in each graph represents median. ^#^*p*<0.05, ^§^*p*<0.01 (Mann-Whitney test).

### Database analyses of Spred2 expression and overall survival

We examined Spred2 expression in bladder cancer database by Sanchez-Carbayo bladder 2 dataset [34], Blaveri bladder 2 [35], and Stransky bladder [36] in a public cancer microarray database (ONCOMINE) [40]. As shown in Figure 6A, Spred2 expression was significantly increased in non-invasive superficial bladder cancer compared to that in normal bladder samples (Fig 6A, left). Of note, Spred2 expression in infiltrating bladder urothelial carcinoma was lower than superficial bladder cancer, which was also found in the other datasets (Fig 6A, middle and right). The decreased Spred2 expression in infiltrating bladder urothelial carcinoma may have affected cancer survival. We then assessed the prognostic value of Spred2 in patients with bladder carcinoma in Kaplan-Meier Plotter (www.kmplot.com). The overall survival for 30 months was higher in patients with higher Spred2 mRNA level (Fig 6B). Although there was no statistical significance in the 150 month-overall survival between the groups (Fig 6C, upper panel), the median survival in Spred2 high expression cohort (42.33 months) was 1.6 times longer than low expression cohort (28.63 months) (Fig 6C, lower panel). Thus, the expression level of Spred2 can be clinically important in the cancer progression.

**Fig 6.**
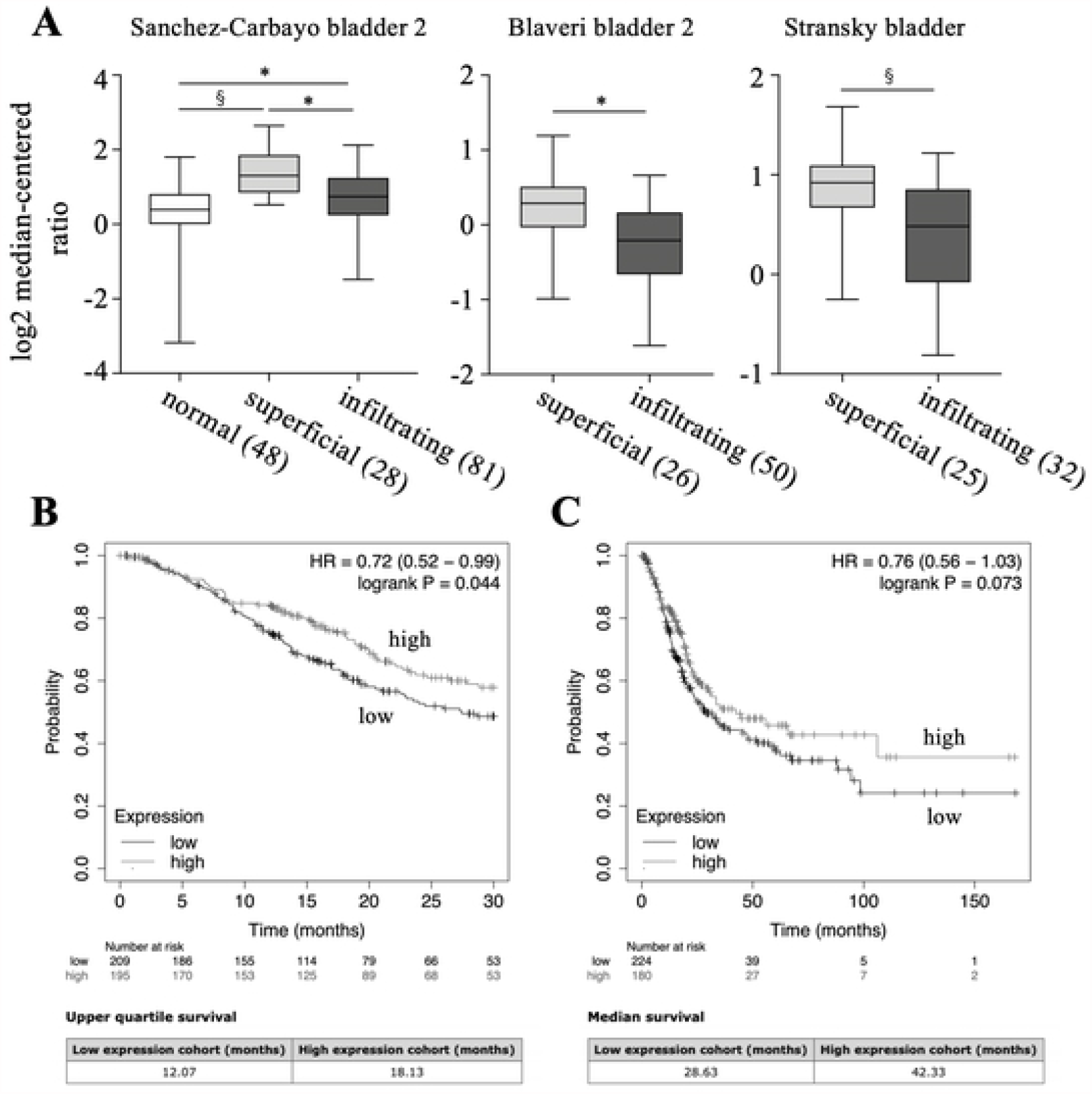
Spred2 expression in overall survival of patients with bladder cancer. (A) Statistical analyses of Spred2 expression in normal, superficial bladder cancer (superficial) and infiltrating bladder urothelial carcinoma (infiltrating) from 3 different datasets (Sanchez-Carbayo bladder 2, Blaveri bladder 2, and Stransky bladder) were shown. The numbers in parentheses indicates the number of samples. ^§^*p*<0.01, ^*^*p*<0.0001 (unpaired two-tailed t test). (B, C) Kaplan-Meier analysis of the data in www.kmplot.com was used to determine the survival probability for 30 months (B) and 150 months (C) of patients with high or low Spred2 expression, followed by the log-rank test.

## Discussion

Cancer cell growth is mediated by various cell signaling pathways. Among them, Ras/Raf/ERK-MAPK is often up-regulated in human diseases including cancer [41], and as such represents an attractive target for the development of anti-cancer drugs [19]. This pathway is also important in urothelial cell migration and invasion [42]. A better understanding of the endogenous negative regulatory mechanism(s) could improve strategies for preventing and treating bladder urothelial tumors. To the best of our knowledge, this is the first report to show Spred2 mRNA and protein expression in bladder urothelial tumors in all categories, ranging from non-tumor to invasive cancer.

Previous studies demonstrated that Spred2 mRNA expression was decreased in hepatocellular carcinoma (HCC) [31] and prostatic adenocarcinoma [32], comparing with that in adjacent non-tumor tissue and benign gland, respectively. Down-regulated Spred2 expression was particularly evident in higher grade prostate cancers [32], and Spred2 expression levels in HCC tissue were inversely correlated with the incidence of tumor invasion and metastasis [31]. These previous findings indicated that Spred2 may function as a potential tumor suppressor gene. In our study, Spred2 mRNA expression was increased in non-invasive cancer HGPUC, whereas the expression in invasive bladder cancer IUC was significantly decreased as compared to that in non-invasive carcinoma HGPUC. Consistently, database analyses showed that Spred2 expression in infiltrating bladder urothelial carcinoma (invasive) was lower than that in superficial bladder cancer (non-invasive). Protein expression of functional Spred2, a membranous positive staining, was frequently observed in LGPUC and HGPUC, but not in CIS and IUC. Thus, Spred2 may play a role as a tumor suppressor in non-invasive carcinomas, but the function appears to be lost in invasive carcinoma.

Spred2 was discovered as a membrane-associated substrates of receptor tyrosine kinases [17, 37]. However; our data indicated that Spred2 was found not only in the membrane but in the cytoplasm in urothelial tumors. Previous confocal microscopy analyses revealed that Spred2 was present in cytoplasm and co-localized with neighbor of BRCA1 (NBR1) [43] or microtubule-associated protein 1A/1B-light chain 3 (*LC3*) [44]. Very interestingly, Spred2-NBR1 complex enhanced Spred2-mediated ERK inhibition upon stimulation with fibroblast growth factor (FGF), suggesting that Spred2/NBR1-dependent down-regulation of ERK-MAPK is achieved via directed endosomal trafficking of activated receptors [43]. Sprouty proteins, a member of the Sprouty/Spred family, were distributed throughout the cytosol, which were underwent rapid translation to membrane ruffles following epidermal growth factor (EGF) stimulation [45]. In urothelial tumors, we showed that membranous Spred2 protein was favorably detected in cancer categories, especially LGPUC and HGPUC. These results indicate that Spred2 may transition from cytoplasm to cell membrane by various stimuli in the cancer microenvironment, exerting the function.

ERK activation was associated with increased Ki67 expression in salivary gland mucoepidermoid carcinoma [39]. Since Spred2 inhibits the ERK pathway and subsequent cell proliferation, we compared the relationship between membranous Spred2 protein expression and pERK score/Ki67 index in each cancer category. Interestingly, HGPUC displaying membranous Spred2 expression showed significantly lower pERK score and Ki67 index, as compared to membrane-negative expression. On the other hand, pERK score was not affected by membranous Spred2 expression in CIS and IUC. Spred2 is presumed to be effective only after reaching a certain level of membrane expression. It appears that ERK activation was so strong in CIS and IUC that concurrent membranous Spred2 expression might be insufficient to suppress the aberrant ERK activation in CIS and IUC. Interestingly, Ki67 index was decreased in IUC with membranous Spred2 expression, although pERK score was not altered by membranous Spred2 expression. Spred2 may downregulate cancer cell proliferation through ERK-MAPK independent pathway in IUC. Spred2 gene mutations can be frequently seen in bladder urothelial carcinoma (Supplementary Fig. 2). The mutated Spred2 may function differently.

Spred2 mRNA expression in CIS was as high as that in HGPUC, however; 75% of CIS showed negative membranous Spred2 staining and only 15% of CIS showed positive membranous Spred2 staining. It remains unclear how Spred2 protein expression is regulated in CIS. The poor correlations were generally reported between the level of mRNA and protein [46, 47]. There are many complicated and varied post-transcriptional mechanisms; post-transcriptional, translational and protein degradation regulation. CIS appears to be the critical turning-point to control the complex regulation. Further study is necessary to understand the specific mechanisms regulating Spred2 mRNA and protein expression.

In conclusion, Spred2 mRNA and protein expression was up-regulated as the grade increased in non-invasive papillary urothelial carcinomas. Membranous Spred2 expression in HGPUC, but not in CIS and IUC, correlated with significantly low levels of ERK activity. In bladder cancer, HGPUC is clinically important because tumor grows more quickly and more likely spread, and tumor progression (invasion) was identified in 40% of all cases [48]. Our present study suggests that Spred2 functions to suppress the growth and progression of cancer in non-invasive bladder cancer through suppressing the ERK pathway, and this regulatory mechanism does not function in invasive bladder cancer.

## Supporting information

### S1 Fig. Immunoreactivity of Spred2 antibody

H1993 cells cultured on Lab-Tek II Slide (8 Chamber, Electron Microscopy Sciences, Hatfield, PA, USA) were transfected with Spred2 expression plasmid (OriGene, Rockville, MD, USA) or control plasmid (OriGene) using turbofectin 8.0 (OriGene). The cells were fixed in 95% ethanol and immunostained with anti-Spred2 polyclonal antibody using the polymer method (Polink-2 Plus HRP RABBIT with DAB kit, GBI, Bothell, WA, USA). Spred2 positive cells were shown in brown.

### S2 Fig. The mutation of Spred2 in cancers

Data were from TCGA Pancancer Atlas from cBioPortal for Cancer Genomics (https://www.cbioportal.org/results/plots). (A) The Spred2 mutations in different cancer. Bladder cancer is the 3rd place having mutation of Spred2 among cancers. (B) The distribution of mutations on the domain structure of Spred2 in bladder urothelial carcinoma.

## Acknowledgments

We thank Mr. Hiroyuki Watanabe and Yasuharu Arashima for their excellent technical assistance. This work was supported in part by JSPS KAKENHI Grant number 25293095, 21H02988 and 16K15258.

## Author contributions

**Conceptualization:** Shinsuke Oda, Akihiro Matsukawa.

**Data curation:** Shinsuke Oda, Akihiro Matsukawa.

**Formal analysis:** Shinsuke Oda, Akihiro Matsukawa.

**Funding acquisition:** Akihiro Matsukawa.

**Investigation:** Shinsuke Oda, Masayoshi Fujisawa, Li Chunning, Takahiro Yamaguchi.

**Methodology:** Shinsuke Oda, Masayoshi Fujisawa, Toshihiro Ito, Teizo Yoshimura.

**Project administration:** Akihiro Matsukawa.

**Supervision:** Akihiro Matsukawa.

**Writing – original draft:** Shinsuke Oda, Masayoshi Fujisawa.

**Writing – review & editing:** Teizo Yoshimura, Toshihiro Ito, Akihiro Matsukawa

